# Genetic equidistance at the nucleotide level

**DOI:** 10.1101/105072

**Authors:** Dejian Yuan, Shi Huang

## Abstract

The genetic equidistance phenomenon was first discovered in 1963 by Margoliash and shows complex taxa to be all approximately equidistant to a less complex species in amino acid percentage identity. The result has been mis-interpretated by the ad hoc universal molecular clock hypothesis, and the much overlooked mystery was finally solved by the maximum genetic diversity hypothesis (MGD). Here, we studied 15 proteomes and their coding DNA sequences (CDS) to see if the equidistance phenomenon also holds at the CDS level. We performed DNA alignments for a total of 5 groups with 3 proteomes per group and found that in all cases the outgroup taxon was equidistant to the two more complex taxa species at the DNA level. Also, when two sister taxa (snake and bird) were compared to human as the outgroup, the more complex taxon bird was closer to human, confirming species complexity rather than time to be the primary determinant of MGD. Finally, we found the fraction of overlap sites where coincident substitutions occur to be inversely correlated with CDS conservation, indicating saturation to be more common in less conserved DNAs. These results establish the genetic equidistance phenomenon to be universal at the DNA level and provide additional evidence for the MGD theory.

## Introduction

The genetic equidistance phenomenon was discovered in the early 1960s when protein alignments among multiple taxa was first performed (1,2). The result of these alignments showed that a low complexity taxon is approximately equidistant to all higher complexity taxa. This was unexpected and immediately led these authors to propose the ad hoc molecular clock hypothesis or similar mutation rates for all taxa. Most researchers failed to realize that this interpretation of the equidistance results is merely a tautology lacking any independent validation (3–6). Many have gone on to try to come up with theories to explain the molecular clock rather than the equidistance (7–12). Among these, the ‘neutral theory’ seems to work the best and has served as the dominant molecular evolutionary theory for half of a century (10–12). However, this theory remains highly controversial and unsupported by hard evidence. According to Kimura and Ohta, the best evidence for the neutral theory is the molecular clock (13). But this is obviously circular reasoning as the neutral theory of molecular evolution was in fact invented because of the molecular clock.

After many years of follow up research, the universal molecular clock hypothesis has now been discredited (14,15). Therefore, the molecular clock and the Neutral theory are clearly false interpretations of the genetic equidistance phenomenon. Ohta’s “nearly neutral theory” explained to some extent some of the problems facing the neutral theory but remains unable to account for the equidistance result (16). Thus, the field still lacks a complete theory as Ohta and Gillespie had acknowledged: “We have yet to find a mechanistic theory of molecular evolution that can readily account for all of the phenomenology.” (17). While genetic diversity within species has long been recognized widely as the chief puzzle of molecular evolution (18,19), the field unfortunately remains unaware of the other key mystery, the genetic equidistance phenomenon, considered by some biologist outside the field as “one of the most astonishing findings of modern science”(20).

The neutral theory treats genetic distance or diversity to be always increasing with time with no limit defined. In contrast, the recently proposed maximum genetic diversity theory (MGD) recognizes two different phases of evolution, linear and saturation (6,21–23). There is ample evidence, including the original equidistance result, to show that sequence divergences are mostly at saturation phase. The saturation level is tightly linked to species biology and organismal complexity (23,24). The MGD of simple organisms is greater than that of complex organisms. The maximum genetic distance between a less complex taxon and all more complex taxa is mainly determined by the MGD of the less complex taxon. The MGD theory includes the proven virtues of the neutral theory as relevant only to microevolution *before* sequence divergence reaches saturation, and explains the equidistance phenomenon as a result of MGD (6,21,22).

Importantly, the equidistance phenomenon has in fact another characteristic, the overlap feature where particular sites have encountered multiple recurrent mutational changes in a particular lineage (25). Overlapped mutant amino acid positions can be easily detected as sites where each taxon has a unique residue when aligning homologous proteins from three different taxa. While the molecular clock may superficially explain the apparent equidistance in numbers, it has not accounted for the overlap feature. The MGD theory explains it as the necessary outcome of mutational saturation. Consistent with being a true interpretation of evolutionary history, the MGD theory has also been instrumental in directing productive research into both evolutionary phylogenetic problems and contemporary biomedical problems (23,25–31).

The genetic equidistance phenomenon has been verified for many proteins and even whole proteomes (5,32,33). However, previous studies only involved amino acid alignments. Here, we examined the phenomenon at the DNA or coding DNA sequence (CDS) level by using whole genome data from multiple taxa.

## Materials and methods

### Whole genome sequences

We downloaded most of the whole genome sequence data used here from Ensembl, which has both peptide sequences and CDS in fasta format. The snake data was downloaded http://gigadb.org/dataset/100196 (34).

### Sequence alignment

We performed multiple groups of three taxa alignments at both the amino acid level and CDS level. Each group of three taxa involved organisms of apparently different complexity with complexity defined by the number of cell types. We only selected taxa whose relative complexity can be easily inferred based on a rough or intuitive estimation of cell type numbers. Most of the alignments were performed by BLAST. We used mkblastdb to build a local BLAST database for both peptide and CDS alignments.

We identified candidate orthologs based on bitscore results of BLASTP by following the reciprocal best hit method (35). The basic procedure entails collecting all the genes in two species and comparing them to one another. If genes from two species identify each other as their closest partners then they are considered orthologs. This works well for closely related species but can be a major problem in highly divergent species. There is a tradeoff between specificity and coverage. For our selection, we restricted the length of the orthologs to be a minimum of 100 amino acids. We aimed for specificity instead of coverage.

### Statistics methods

To calculate the average protein identity between two proteomes, longer proteins should be weighted more. Thus, to adjust the effect of protein length in amino acid alignments, we used the formula (identity x length)/average_length to obtain a weighted protein identity score (33). Other methods include t test, f test, and Spearman or Pearson correlation analysis.

## Calculation of overlap ratio

The overlap ratio is defined as the number of actual overlap positions divided by the number of candidate positions (25). The candidate overlap positions in any three taxa comparison involving an outgroup include all the different positions between the two sister lineages. We used ClustalW to create a three taxa alignment document of either peptides or CDS. We then used a custom script to count the number of overlap positions where each of the three species has a unique residue. For CDS comparison, an overlap site was defined as a nucleotide position where each of the three aligned taxa had a unique nucleotide.

## Results

### Genetic equidistance at both amino acid and CDS levels

We first performed peptide alignments for a selected set of proteomes. To study genetic equidistance, a minimum of three species is needed where two complex taxa could be compared to a less complex taxon to determine whether they are equidistance to the less complex taxon. Complexity is inferred based on an intuitive estimation of cell type numbers. We only selected those taxa where difference in complexity is intuitively obvious. We selected the following 15 taxa: Homo sapiens, Xenopus tropicalis (frog), Oreochromis niloticus (fish), Pan troglodytes (chimpanzee), Equus caballus (horse), Sus scrofa (wild pig), Gallus gallus (chicken), Pongo abelii (orangutan), Felis catus (cat), nolis carolinensis (lizard), Ailuropoda melanoleuca (panda), Canis familiaris (dog), Pelodiscus sinensis (turtle), Deinagkistrodon acutus (snake), Cuculus canorus (bird). From these taxa, we designed 6 sets of three taxa combinations where the three taxa are of different apparent complexity. For 5 of these sets, the outgroup was the least complex taxon (Table 1). For the remaining set, the outgroup was human while the sister taxa were snake and bird (Table 1).

**Table 1.** Protein alignments

| Species |  |  | Average percentage identity |  |  |  | Average length |  |  | Average weighted identity |  |  |  | Protein number |
| --- | --- | --- | --- | --- | --- | --- | --- | --- | --- | --- | --- | --- | --- | --- |
| 1st | 2nd | 3rd | 1-2 | 2-3 | 1-3 | $P^1$ | 1-2 | 2-3 | 1-3 | 1-2 | 2-3 | 1-3 | $P^1$ | |
| HOMO | XENTR | ORENI | 69.75 | 64.62 | 64.93 | 0.1903 | 577.3 | 557.9 | 579.6 | 68.90 | 63.44 | 64.03 | 0.540 | 8023 |
| SUSSC | XENTR | ORENI | 68.98 | 64.32 | 64.35 | 0.9046 | 529.5 | 552.6 | 529.3 | 67.95 | 63.13 | 63.36 | 0.798 | 7255 |
| HOMO | PANTR | EQUCA | 98.74 | 88.61 | 89.00 | <0.001 | 585.6 | 549.6 | 560.2 | 98.74 | 88.97 | 89.35 | 0.712 | 12642 |
| PONAB | FELCA | ANOCA | 88.91 | 71.45 | 71.10 | 0.1069 | 583.0 | 558.2 | 554.1 | 88.61 | 70.99 | 70.73 | 0.767 | 10640 |
| ALIME | CANFA | PELSI | 93.73 | 74.46 | 74.36 | 0.636 | 585.7 | 548.3 | 544.1 | 93.66 | 74.31 | 74.24 | 0.940 | 10507 |
| HOMO | CUCCA | DEIAC | 76.50 | 77.65 | 73.05 | <0.001 | 549.4 | 520.1 | 531.6 | 75.87 | 76.96 | 72.32 | <0.001 | 5940 |

Notes: <sup>1</sup> $P$ values were t test results between 1<sup>st</sup>-3<sup>rd</sup> and 2<sup>nd</sup>-3<sup>rd</sup> comparisons.

Notes: ^1^*P* values were t test results between 1^st^–3^rd^ and 2^nd^–3^nd^ comparisons.

HOMO：: Homo sapiens
XENTR：: Xenopus tropicalis (frog)
ORENI：: Oreochromis niloticus (Nile tilapia fish)
PANTR：: Pan troglodytes (chimpanzee)
EQUCA：: Equus caballus (horse)
SUSSC：: Sus scrofa (wild pig)
GALGA：: Gallus gallus (chicken)
PONAB：: Pongo abelii (orangutan)
FELCA：: Felis catus (cat)
ANOCA：: Anolis carolinensis (lizard)
ALIME：: Ailuropoda melanoleuca (giant panda)
CANFA：: Canis familiaris (dog)
PELSI：: Pelodiscus sinensis (Chinese softshell turtle)
DEIAC：: Deinagkistrodon acutus (snake)
CUCCA：: Cuculus canorus (bird)

We selected orthologs of at least 100 amino acids in length for sequence comparison. For each of the 6 sets, the average alignment gaps between each pair of orthologs within the set were similar (Supplementary Table S1-S6). We obtained the percentage identity for each protein. We also calculated a weighted identity score for each protein by the formula: (percent identity x protein length) / average length of all proteins. The weighted score is more suitable for a meaningful average protein identity as longer length proteins should contribute more to the proteome average than short ones. The average pairwise identity and weighted identity scores among the orthologs in each set are shown in Table 1 and Supplementary Table S1-S6.

All of the 5 groups of comparisons involving a less complex outgroup showed the expected results consistent with the MGD theory, including human and frog being equidistant to fish, pig and frog being equidistant to fish, human and chimpanzee being equidistant to horse (here only the weighted protein identity showed the expected result), orangutan and cat being equidistant to lizard, and panda and dog being equidistant to turtle (Table 1).

To further examine the role of complexity versus time factor in the equidistance phenomenon, we compared human, bird, and snake where human serves as the more complex outgroup that is equidistant to bird and snake in terms of evolutionary time. One would expect to see human to be equidistant to bird and snake if the molecular clock is real. On the other hand, one would expect human to be closer to bird than to snake if complexity is the main determinant of MGD and since bird may be more complex than snake. The result showed that human is closer to bird than to snake (*P* < 0.00001), thus supporting the MGD prediction (Table 1). Bird was found to be closer to snake than to human, which may reflect shared physiology as oviparous animals.

We then performed the corresponding CDS alignments for the above 6 groups and obtained similar results (Table 2). The number of homologous proteins used for the CDS alignments was less than those for the protein alignments because we only retained CDS orthologs that showed reciprocal best hits for both amino acid and CDS alignments.

**Table 2.** CDS alignments.

| Species |  |  | Average percentage identity |  |  |  | Average length |  |  | CDS number |
| --- | --- | --- | --- | --- | --- | --- | --- | --- | --- | --- |
| 1st | 2nd | 3rd | 1-2 | 2-3 | 1-3 | $P^1$ | 1-2 | 2-3 | 1-3 | |
| HOMO | XENTR | ORENI | 80.70 | 78.43 | 78.90 | 0.303 | 2182 | 1586 | 1958 | 103 |
| SUSSC | XENTR | ORENI | 81.20 | 78.92 | 79.26 | 0.547 | 1611 | 1502 | 1592 | 84 |
| HOMO | PANTR | EQUCA | 99.35 | 90.41 | 90.42 | 0.699 | 1729 | 1570 | 1628 | 12363 |
| PONAB | FELCA | ANOCA | 92.79 | 81.51 | 81.69 | 0.140 | 2088 | 1633 | 1618 | 2210 |
| ALIME | CANFA | PELSI | 95.43 | 82.42 | 82.40 | 0.857 | 2137 | 1694 | 1668 | 3587 |
| HOMO | CUCCA | DEIAC | 83.39 | 83.57 | 81.72 | <0.01 | 1850 | 1908 | 1954 | 916 |

Notes: <sup>1</sup> $P$ values were t test results between 1<sup>st</sup>-3<sup>rd</sup> and 2<sup>nd</sup>-3<sup>rd</sup> comparisons.

Notes: 1*P* values were t test results between 1^st^–3^rd^ and 2^nd^–3^nd^ comparisons.

We next analyzed the overlap sites at CDS level for the human, frog, and fish group involving a total of 8201 genes (Supplementary Table S7). We counted the number of overlapped and candidate positions and calculated the overlap ratio as the number of overlapped nucleotide positions divided by the number of candidate nucleotide positions. The results showed a significant inverse correlation between the overlap ratio and CDS conservation (R=−0.63, *P* <2.20E-16, Figure 1), consistent with saturation being the reason for the overlap as proposed by the MGD theory.

**Figure 1.**
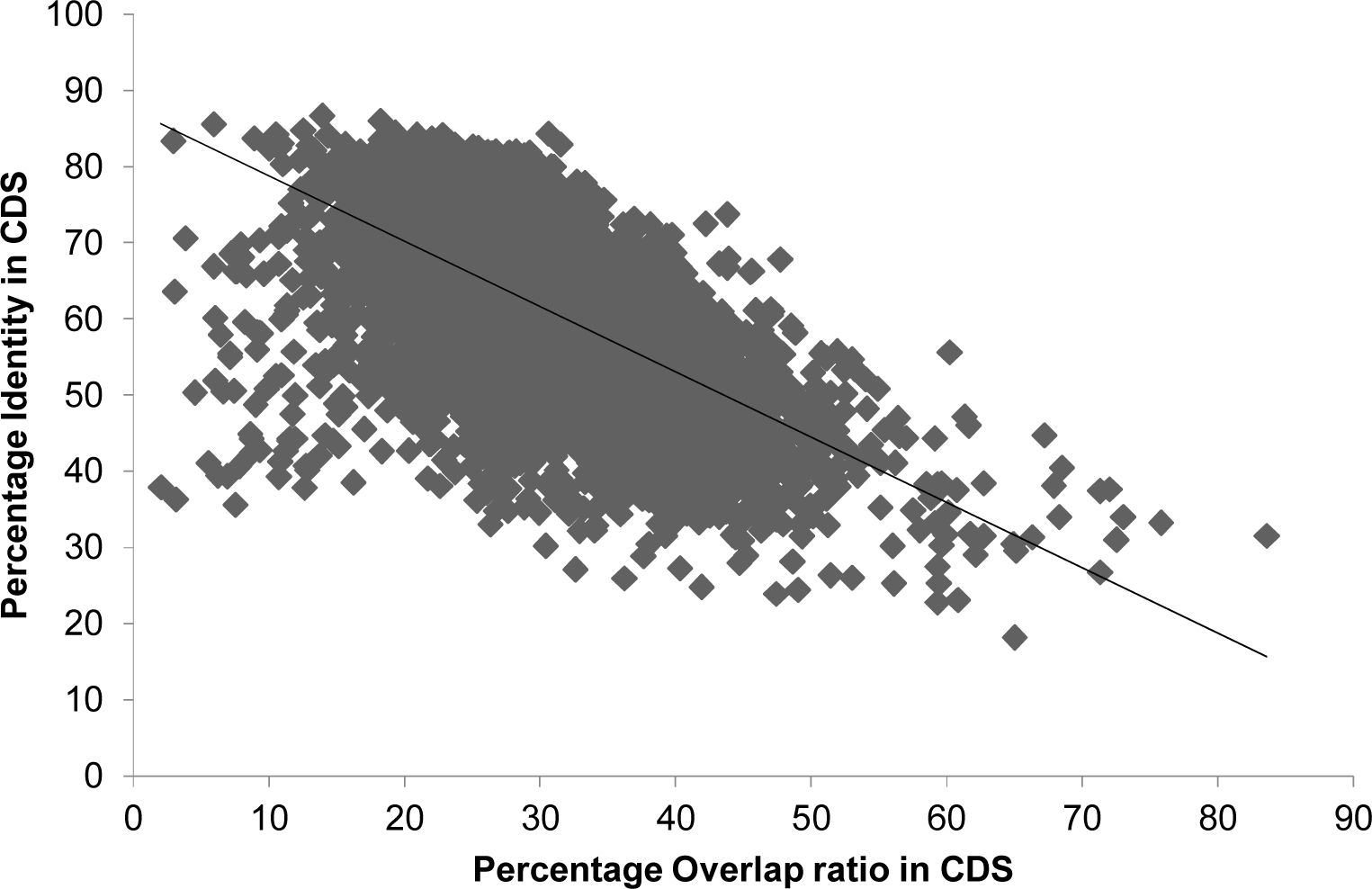
**The overlap ratio in CDS alignments**. CDS alignments were performed for human, frog, and fish involving a total of 8201 genes. The overlap ratio was determined for each gene and plotted against the gene’s percentage identity between human and frog.

## Discussion

Our analysis here shows that the genetic equidistance phenomenon also holds at the genome wide DNA or CDS level, further establishing its universal nature. As our analysis involved a large number of taxa selected for no better reason than the availability of whole genome wide CDS sequences, the results indicate the universality of genetic equidistance with regard to species. The universal nature of this phenomenon in terms of both proteomes and CDS suggests that it is a direct outcome of mutation events at the DNA level. It is further inferred that the equidistance phenomenon should also hold in non CDS regions, but low conservation may make it technically difficult to find orthologs required for a meaningful comparison. Consistent with the expectation of equidistance in non-CDS regions, it has been recently shown that Africans are equidistant to non-Africans as measured by genome wide SNPs, which are mostly non-coding (36). Such equidistance is maximum distance because genome wide SNP amounts are at saturation level as indicated by more sharing of faster evolving SNPs among different human groups (36), and higher SNP amounts in patient populations with complex diseases (27).

Our finding here that human is closer to birds than to snakes in both proteomes and CDS confirms the prediction by the MGD theory and invalidates the molecular clock and the neutral theory. MGD is largely determined by physiology or species complexity as suggested by the MGD theory. While one would expect snakes to be equidistant to humans and birds according to the MGD theory, our results showed closer distance between snakes and birds relative to snakes and humans. This is likely due to significant physiological differences between oviparous animals and placenta mammals. As MGD is tightly linked to physiology, the shared physiology between snakes and birds may prohibit certain genetic variations that would be tolerated in other animals.

There are two kinds of equidistance, linear and maximum (22). For short evolutionary time scales or for slow-evolving sequences, one would observe the linear genetic equidistance phenomenon where the molecular clock or the neutral theory holds and the distance is nearly linearly related to time. Over long evolutionary time scales or for fast-evolving sequences, maximum genetic equidistance would be observed and most parts of any genome today are at the saturation phase of evolution. These two kinds of equidistance can be easily distinguished by the overlap feature (25). Our results on the overlap ratio at the DNA level here further established the inverse relationship between the overlap ratio and DNA conservation. Any future phylogenetic studies must pay attention to the overlap feature or the saturation phenomenon and should only select and use slow evolving genes that show the least amount of overlap sites (31).

## Acknowledgements

This work was supported by the National Natural Science Foundation of China (Grant No. 81171880) and the National Basic Research Program of China (Grant No. 2011CB51001).

